# Sodium Thiosulfate acts as an H_2_S mimetic to prevent intimal hyperplasia via inhibition of tubulin polymerization

**DOI:** 10.1101/2021.09.09.459573

**Authors:** Diane Macabrey, Alban Longchamp, Michael R. MacArthur, Martine Lambelet, Severine Urfer, Jean-Marc Corpataux, Sebastien Deglise, Florent Allagnat

**Affiliations:** Department of Vascular Surgery, Lausanne University Hospital, Switzerland; Department of Biomedical Sciences, University of Lausanne, Switzerland; Department of Health Sciences and Technology, Swiss Federal Institute of Technology (ETH) Zurich, Zurich, Switzerland

**Keywords:** Intimal hyperplasia, smooth muscle cells, proliferation, hydrogen sulfide, sodium thiosulfate

## Abstract

**Background:** Intimal hyperplasia (IH) remains a major limitation in the long-term success of any type of revascularization. IH is due to vascular smooth muscle cell (VSMC) dedifferentiation, proliferation and migration. The gasotransmitter Hydrogen Sulfide (H_2_S) inhibits IH in pre-clinical models. However, there is currently no clinically approved H_2_S donor. Here we used sodium thiosulfate (STS), a clinically-approved source of sulfur, to limit IH.

**Methods:** Hypercholesterolemic LDLR deleted (LDLR^-/-^), WT or CSE^-/-^ male mice randomly treated with 4g/L STS in the water bottle were submitted to focal carotid artery stenosis to induce IH. Human vein segments were maintained in culture for 7 days to induce IH. Further *in vitro* studies were conducted in primary human vascular smooth muscle cell (VSMC).

**Findings:** STS inhibited IH in mice and in human vein segments. STS inhibited cell proliferation in the carotid artery wall and in human vein segments. STS increased polysulfides *in vivo* and protein persulfidation *in vitro*, which correlated with microtubule depolymerization, cell cycle arrest and reduced VSMC migration and proliferation.

**Interpretation:** STS, a drug used for the treatment of cyanide poisoning and calciphylaxis, protects against IH in a mouse model of arterial restenosis and in human vein segments. STS acts as an H_2_S donor to limit VSMC migration and proliferation via microtubule depolymerization.

**Funding:** This work was supported by the Swiss National Science Foundation (grant FN-310030_176158 to FA and SD and PZ00P3-185927 to AL); the Novartis Foundation to FA; and the Union des Sociétés Suisses des Maladies Vasculaires to SD.

**Graphical Abstract:** 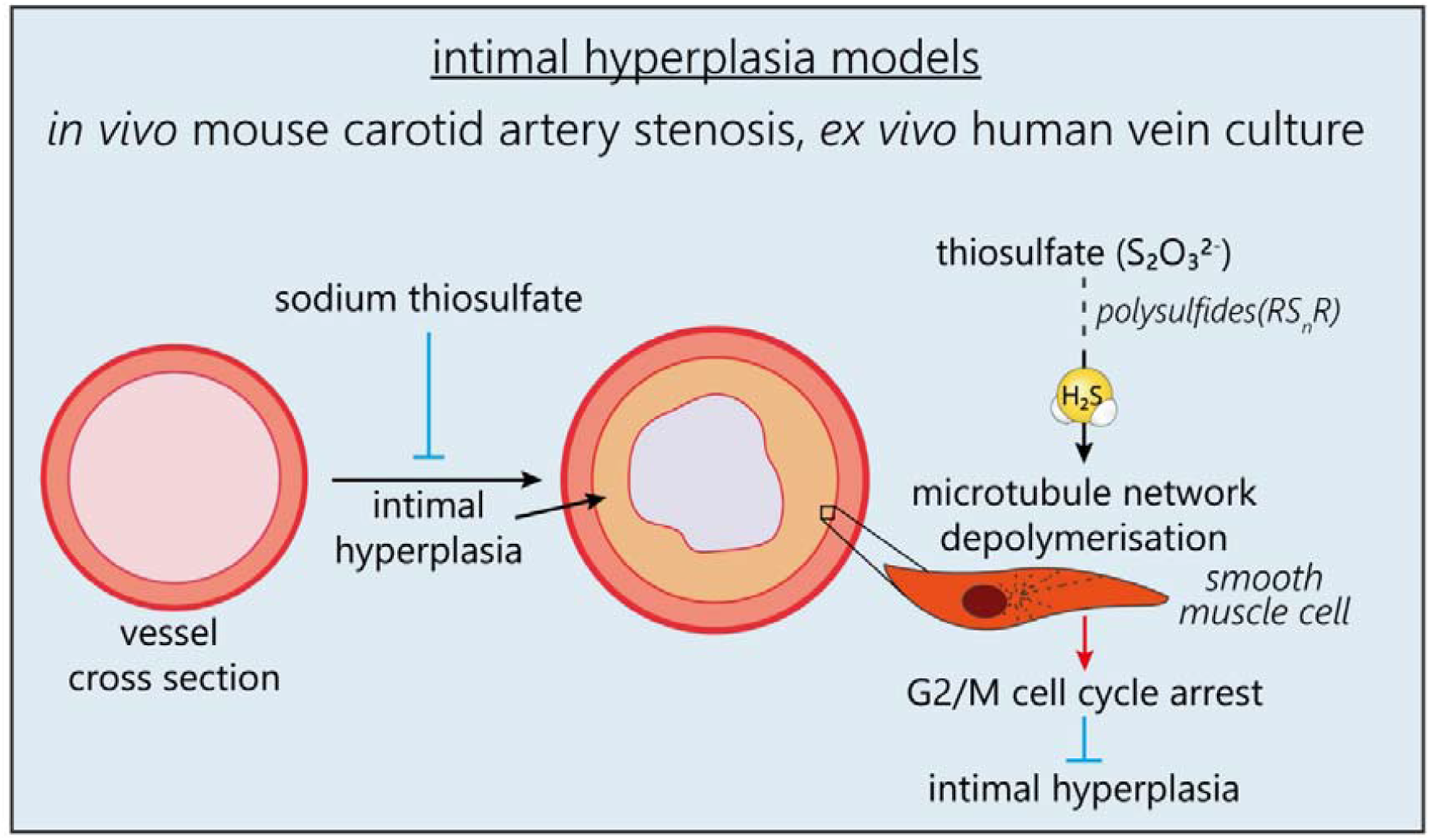

**Research in context:** *Evidence before this study:* Intimal hyperplasia (IH) is a complex process leading to vessel restenosis, a major complication following cardiovascular surgeries and angioplasties. Therapies to limit IH are currently limited. Pre-clinical studies suggest that hydrogen sulfide (H_2_S), an endogenous gasotransmitter, limits restenosis. However, despite these potent cardiovascular benefits in pre-clinical studies, H_2_S-based therapeutics are not available yet. Sodium thiosulfate (Na_2_S_2_O_3_) is an FDA-approved drug used for the treatment of cyanide poisoning and calciphylaxis, a rare condition of vascular calcification affecting patients with end-stage renal disease. Evidence suggest that thiosulfate may generate H_2_S *in vivo* in pre-clinical studies.

*Added value of this study:* Here, we demonstrate that STS inhibit IH in a surgical mouse model of IH and in an *ex vivo* model of IH in human vein culture. We further found that STS increases circulating polysulfide levels *in vivo* and inhibits IH via decreased cell proliferation via disruption of the normal cell’s cytoskeleton. Finally, using CSE knockout mice, the main enzyme responsible for H_2_S production in the vasculature, we found that STS rescue these mice from accelerated IF formation.

*Implications of all the available evidence:* These findings suggest that STS holds strong translational potentials to limit IH following vascular surgeries and should be investigated further.

## 1. Introduction

Prevalence of peripheral arterial disease continues to rise worldwide, largely due to the combination of aging, smoking, hypertension, and diabetes mellitus (1-3). Vascular surgery, open or endo, remains the best treatment. However, the vascular trauma associated with the intervention eventually lead to restenosis and secondary occlusion of the injured vessel. Re-occlusive lesions result in costly and complex recurrent end-organ ischemia, and often lead to loss of limb, brain function, or life. Despite the advent of new medical devices such as drug eluting stent and drug-coated balloons, restenosis has been delayed rather than suppressed (4), and limited therapies are currently available.

Restenosis is mainly due to intimal hyperplasia (IH), a process instated by endothelial cell injury and inflammation, which induces vascular smooth muscle cell (VSMC) reprogramming. VSMCs become proliferative and migrating, secrete extra-cellular matrix and form a new layer called the neo-intima, which slowly reduces the vessel luminal diameter (5).

Hydrogen sulfide (H_2_S) is an endogenously produced gasotransmitter derived from cysteine metabolism (6). Circulating H_2_S levels are reduced in humans suffering from vascular occlusive disease (7, 8) and pre-clinical studies using water-soluble sulfide salts such as Na_2_S and NaHS have shown that H_2_S has cardiovascular protective properties(6), including reduction of IH *in vivo* in rats (9), rabbits (10), mice (11), and *ex-vivo* in human vein segments (12). However, the fast and uncontrolled release, narrow therapeutic range and high salt concentration of these compounds limit their therapeutic potential. Due to these limitations, H_2_S-based therapy are currently not available.

Here, we focused on sodium thiosulfate (Na_2_S_2_O_3_), an FDA-approved drug used in gram-quantity doses for the treatment of cyanide poisoning(13) and calciphylaxis, a rare condition of vascular calcification affecting patients with end-stage renal disease(14). Pharmaceutical-grade sodium thiosulfate (STS) is available and has been suggested to release H_2_S through non-enzymatic and enzymatic mechanisms (15, 16).

We tested whether STS inhibit IH in a surgical mouse model of IH and in an *ex vivo* model of IH in human vein culture. NaHS, a validated H2S donor was systematically compared to STS. We observed that STS was at least as potent as NaHS to inhibit IH in mice carotids, and in human vein. Although STS did not release detectable amounts of H2S or polysulfides *in vitro*, it increased protein persulfidation and circulating polysulfide levels *in vivo*. STS inhibited apoptosis and matrix deposition associated with the development of IH, as well as VSMC proliferation and migration. We further observed that STS and NaHS induced microtubule depolymerization in VSMCs.

## 2. Methods

For details on materials and reagents please see the Supplementary Material files

### 2.1. Mouse treatment

8 to 10 weeks old male WT or LDL receptor knock out (LDLR^-/-^) mice on a C57BL/6J genetic background were randomly divided into control vs sodium thiosulfate (STS) or NaHS. Sodium Thiosulfate (Hänseler AG, Herisau, Switzerland) was given in mice water bottle at 4g/L (0.5g/Kg/day), changed 3 times a week. NaHS (Sigma-Aldrich) was given in mice water bottle at 0.5g/L (125mg/Kg/day), changed every day. LDLR^-/-^ mice were put on a cholesterol rich diet (Western 1635, 0.2% Cholesterol, 21% Butter, U8958 Version 35, SAFE^®^ Complete Care Competence) for 3 weeks prior to experiments initiation. Mice were euthanized after 7 or 28 days of treatment by cervical dislocation under isoflurane anesthesia (inhalation 3% isoflurane under 2.5 liter of O_2_) followed by PBS perfusion. Aortas, carotid arteries, livers and serum or plasma (via intracardiac blood collection with a 24G needle) were harvested.

### 2.2. Cse^-/-^ mice

Cse knockout mice were generated from a novel floxed line generated by embryonic injection of ES cells containing a Cth allele with LoxP sites flanking exon 2 (Cth tm1a(EUCOMM)Hmgu). Both ES cells and recipient embryos were on C57BL/6J background. Mice that were homozygous for the floxed allele were crossed with CMV-cre global cre-expressing mice (B6.C-Tg(CMV-cre)1Cgn/J), which have been backcrossed with C57BL/6J for 10 generations to create constitutive whole-body Cse^-/-^ animals on a C57BL/6J background. The line was subsequently maintained by breeding animals heterozygous for the Cse null allele. Mouse ear biopsies were taken and digested in DirectPCR lysis reagent (Viagen Biotech, 102-T) with proteinase K (Qiagen, 1122470). WT, heterozygous and knockout mice were identified by PCR using the forward primer 5’-AGC ATG CTG AGG AAT TTG TGC-3’ and reverse primer 5’-AGT CTG GGG TTG GAG GAA AAA-3’ to detect the WT allele and the forward primer 5’-TTC AAC ATC AGC CGC TAC AG-3’ to detect knock-out allele using the platinum Taq DNA Polymerase (Invitrogen, cat#10966-026)

### 2.3. Carotid artery stenosis (CAS) surgery

The carotid artery stenosis (CAS) was performed as previously published (17) on 8 to 10 week old male WT or LDLR^-/-^ mice. Treatment was initiated 3 days before surgery and continued for 28 days post-surgery until organ collection. The day of the surgery, mice were anesthetized with an intraperitoneal injection of Ketamine (80mg/kg) (Ketasol-100, Gräub E.Dr.AG, Bern Switzerland) and Xylazine (15 mg/kg) (Rompun®, Provet AG, Lyssach, Switzerland). The left carotid artery was exposed and separated from the jugular vein and vagus nerve. Then, a 7.0 PERMA silk (Johnson & Johnson AG, Ethicon, Zug, Switzerland) thread was looped and tightened around the carotid in presence of a 35-gauge needle. The needle was removed, thereby restoring blood flow, albeit leaving a significant stenosis. The stenosis triggers IH proximal to the site of injury, which was measured 28 days post surgery (17). Buprenorphine (0.05 mg/kg Temgesic, Reckitt Benckiser AG, Switzerland) was provided subcutaneously as post-operative analgesic every 12 hours for 24 hours. Mice were euthanized under isoflurane anesthesia (inhalation 3% isoflurane under 2.5 liter of O_2_) by cervical dislocation and exsanguination, perfused with PBS followed by buffered formalin 4% through the left ventricle. Surgeons were blind to the group during surgeries.

### 2.4. Human tissue and VSMC culture

Static vein culture was performed as previously described (12, 18). Briefly, the vein was cut in 5 mm segments randomly distributed between conditions. One segment (D0) was immediately preserved in formalin or flash frozen in liquid nitrogen and the others were maintained in culture for 7 days in RPMI-1640 Glutamax I supplemented with 10 % FBS and 1% antibiotic solution (10,000 U/mL penicillin G, 10,000 U/mL streptomycin sulfate) in cell culture incubator at 37°C, 5% CO_2_ and 21% O_2_.

Human VSMCs were also prepared from these human great saphenous vein segments as previously described (12, 18). Vein explants were plated on the dry surface of a cell culture plate coated with 1% Gelatin type B (Sigma-Aldrich). Explants were maintained in RPMI, 10% FBS medium in a cell culture incubator at 37°C, 5% CO_2_, 5% O_2_ environment.

### 2.5. Carotid and human vein histomorphometry

After 7 days in culture, or immediately upon vein collection (D0), the vein segments were fixed in buffered formalin, embedded in paraffin, cut into 5 µm sections, and stained using Van Gieson Elastic Laminae (VGEL) staining as previously described (12, 19). Three photographs per section were taken at 100x magnification and 8 measurements of the intima and media thicknesses were made, evenly distributed along the length of the vein wall.

Left ligated carotids were isolated and paraffin-embedded. Six-µm sections of the ligated carotid artery were cut from the ligature towards the aortic arch and stained with VGEL for morphometric analysis. Cross sections at every 300 µm and up to 2 mm from the ligature were analyzed using the Olympus Stream Start 2.3 software (Olympus, Switzerland). For intimal and medial thickness, 72 (12 measurements/cross section on six cross sections) measurements were performed, as previously described (12).

Two independent researchers blind to the experimental groups did the morphometric measurements, using the Olympus Stream Start 2.3 software (Olympus, Switzerland) (12).

### 2.6. H_2_S and polysulfide measurement

Free H_2_S was measured in cells using the SF_7_-AM fluorescent probe (20) (Sigma-Aldrich). The probe was dissolved in anhydrous DMF at 5 mM and used at 5 μM in serum-free RPMI medium with or without VSMCs. Free polysulfide was measured in cells using the SSP4 fluorescent probe. The probe was dissolved in DMF at 10 mM and diluted at 10 μM in serum-free RPMI medium with or without VSMCs. Fluorescence intensity (λ_ex_ = 495 nm; λ_em_ = 520 nm) was measured continuously in a Synergy Mx fluorescent plate reader (BioTek Instruments AG, Switzerland) at 37 °C before and after addition of various donors, as indicated.

Plasma polysulfides were measured using the SSP4 fluorescent probe. Plasma samples were diluted 3 times and incubated for 10 min at 37°C in presence of 10 μM SSP4. Plasma polysulfides were calculated using a Na_2_S_3_ standard curve. Liver polysulfides were measured using the SSP4 fluorescent probe. Pulverized frozen liver was resuspended in PBS-0.5% triton X-100, sonicated and adjusted to 0.5 mg/ml protein concentration. Lysates were incubated for 15 min at 37°C in presence of 10 μM SSP4 and fluorescence intensity (λ_ex_ = 495 nm; λ_em_ = 520 nm) was measured in a Synergy Mx fluorescent plate reader (BioTek Instruments AG, Switzerland)

### 2.7. Persulfidation protocol

Persulfidation protocol was performed using a dimedone-based probe as recently described (21). Persulfidation staining was performed on VSMCs grown on glass coverslips. Briefly, 1mM 4-Chloro-7-nitrobenzofurazan (NBF-Cl, Sigma) was diluted in PBS and added to live cells for 20 minutes. Cells were washed with PBS then fixed for 10 minutes in ice-cold methanol. Coverslips were rehydrated in PBS, and incubated with 1mM NBF-Cl for 1h at 37°C. Daz2-Cy5.5 (prepared with 1mM Daz-2, 1mM alkyne Cy5.5, 2mM copper(II)-TBTA, 4mM ascorbic acid with overnight incubation at RT, followed by quenching for 1h with 20mM EDTA) was added to the coverslips and incubated at 37°C for 1h. After washing with methanol and PBS, coverslips were mounted in Vectashield mounting medium with DAPI and visualized with a 90i Nikon fluorescence microscope.

### 2.8. BrdU assay

VSMC were grown at 80% confluence (5·10^3^ cells per well) on glass coverslips in a 24-well plate and starved overnight in serum-free medium. Then, VSMC were either treated or not (ctrl) with the drug of choice for 24 hours in full medium (RPMI 10% FBS) in presence of 10µM BrdU. All conditions were tested in parallel. All cells were fixed in ice-cold methanol 100% after 24 hours of incubation and immunostained for BrdU. Images were acquired using a Nikon Eclipse 90i microscope. BrdU-positive nuclei and total DAPI-positive nuclei were automatically detected using the ImageJ software (12).

### 2.9. Flow Cytometry

VSMC were grown at 70% confluence (5·10^4^ cells per well) and treated for 48 hours with 15mM STS or 10nM Nocodazole. Then, cells were tripsinized, collected and washed in ice-cold PBS before fixation by dropwise addition of ice-cold 70% ethanol while slowly vortexing the cell pellet. Cells were fixed for 1 hour at 4°C, washed 3 times in ice-cold PBS and resuspended in PBS supplemented with 20µg/mL RNAse A and 10µg/mL DAPI. Flow cytometry was performed in a Cytoflex-S apparatus (Beckmann).

### 2.10. Wound healing assay

VSMC were grown at confluence (10^4^ cells per well) in a 12-well plate and starved overnight in serum-free medium. Then, a scratch wound was created using a sterile p200 pipette tip and medium was changed to full medium (RPMI 10% FBS). Repopulation of the wounded areas was recorded by phase-contrast microscopy over 24 hours in a Nikon Ti2-E live cell microscope. All conditions were tested in parallel. The area of the denuded area was measured at t=0 hours and t=10 hours after the wound using the ImageJ software by two independent observers blind to the conditions. Data were expressed as a ratio of the healed area over the initial wound area.

### 2.11. Apoptosis assay

Apoptosis TUNEL assay was performed using the DeadEnd™ Fluorometric TUNEL system kit (Promega) on frozen sections of human vein segments. Immunofluorescent staining was performed according to the manufacturer’s instruction. Apoptotic nuclei were automatically detected using the ImageJ software and normalized to the total number of DAPI-positive nuclei. *In vitro* VSMC apoptosis was determined by hoescht/propidium iodide staining of live VSMC and manually counted by two independent blinded experimenters (22).

### 2.12. Immunohistochemistry

#### Polychrome Herovici staining

was performed on paraffin sections as described (23). Young collagen is stained blue, while mature collagen is pink. Cytoplasm is counterstained yellow. Hematoxylin is used to counterstain nuclei blue to black.

#### Collagen III staining

was performed on frozen sections (OCT embedded) of human vein segments using mouse anti-Collagen III antibody (ab7778, abcam). Briefly, tissue slides were permeabilized in PBS supplemented with 2 wt. % BSA and 0.1 vol. % Triton X-100 for 30 min, blocked in PBS supplemented with 2 wt. % BSA and 0.1 vol. % Tween 20 for another 30 min, and incubated overnight with the primary antibodies diluted in the same buffer. The slides were then washed 3 times for 5 min in PBS supplemented with 0.1 vol. % Tween 20, and incubated for 1 hour at room temperature with anti-mouse AlexaFluor 568 (1/1000, ThermoFischer). Slides were visualized using a Nikon 90i fluorescence microscope (Nikon AG). Collagen III immunostaining area was quantified using the ImageJ software and normalized to the total area of the vein segment.

#### PCNA (proliferating cell nuclear antigen) and α-tubulin immunohistochemistry

was performed on paraffin sections as previously described (24). After rehydration and antigen retrieval (TRIS-EDTA buffer, pH 9, 17 min in a microwave at 500 watts), immunostaining was performed on human vein or carotid sections using the EnVision +/HRP, DAB+ system according to manufacturer’s instructions (Dako, Baar, Switzerland). Slides were further counterstained with hematoxylin. PCNA and hematoxylin positive nuclei were manually counted by two independent observers blinded to the conditions.

#### α-tubulin immunofluorescent staining

in human VSMC was performed as previously described. Cell were fixed at -20°C for 10min in absolut methanol. Then, cell were blocked/permeabilized in PBS-triton 0.2%, BSA 3% for 45 min at room temperature. Cells were incubated overnight at 4°C in the primary antibody diluted in PBS-0.1% tween, 3% BSA, washed 3 times in PBS and incubated for 1 hour at room temperature with the secondary antibody diluted in PBS-0.1% tween, 3% BSA, washed again 3 times in PBS and mounted using Vectashield mounting medium for fluorescence with DAPI. The microtubule staining was quantified automatically using FiJi (ImageJ, 1.53c). Image processing was as follows: Plugin, Tubeness/Process, Make Binary/Analyze, Skeleton. Data were summarized as filament number and total length, normalized to the number of cells per images. Data were generated from images from 3 independent experiments, 3 to 4 images per experiment per condition.

### 2.13. Western blotting

Mice aortas or human vein segments were flash-frozen in liquid nitrogen, grinded to power and resuspended in SDS lysis buffer (62.5 mM TRIS pH6,8, 5% SDS, 10 mM EDTA). Protein concentration was determined by DC protein assay (Bio-Rad Laboratories, Reinach, Switzerland). 10 to 20 µg of protein were loaded per well. Primary cells were washed once with ice-cold PBS and directly lysed with Laemmli buffer as previously described (12, 18). Lysates were resolved by SDS-PAGE and transferred to a PVDF membrane (Immobilon-P, Millipore AG, Switzerland). Immunoblot analyses were performed as previously described(18) using the antibodies described in supplementary Table S1. Membranes were revealed by enhanced chemiluminescence (Immobilon, Millipore) using the Azure Biosystems 280 and analyzed using Image J. Protein abundance was normalized to total protein using Pierce™ Reversible Protein Stain Kit for PVDF Membranes.

### 2.14. In vitro tubulin polymerization assay

The assay was performed using the *In Vitro* Tubulin Polymerization Assay Kit (≥99% Pure Bovine Tubulin; 17-10194 Sigma-Aldrich), according to the manufacturer’s instruction.

### 2.15. Statistical analyses

All experiments adhered to the ARRIVE guidelines and followed strict randomization. A power analysis was performed prior to the study to estimate sample-size. We hypothesized that STS would reduce IH by 50%. Using an SD at +/- 20% for the surgery and considering a power at 0.9, we calculated that n= 12 animals/group was necessary to validate a significant effect of the STS. All experiments were analyzed using GraphPad Prism 8. One or 2-ways ANOVA were performed followed by multiple comparisons using post-hoc t-tests with the appropriate correction for multiple comparisons.

### 2.17. Ethics Statement

Human great saphenous veins were obtained from donors who underwent lower limb bypass surgery (25). Written, informed consent was obtained from all vein donors for human vein and VSMC primary cultures. The study protocols for organ collection and use were reviewed and approved by the Centre Hospitalier Universitaire Vaudois (CHUV) and the Cantonal Human Research Ethics Committee (http://www.cer-vd.ch/, no IRB number, Protocol Number 170/02), and are in accordance with the principles outlined in the Declaration of Helsinki of 1975, as revised in 1983 for the use of human tissues.

All animal experimentations conformed to *the National Research Council:* Guide for the Care and Use of Laboratory Animals (26). All animal care, surgery, and euthanasia procedures were approved by the CHUV and the Cantonal Veterinary Office (SCAV-EXPANIM, authorization number 3114, 3258 and 3504).

## 3. Results

### 3.1. STS limits IH development in mice after carotid artery stenosis

We first assessed whether STS protects against IH, 28 days after carotid artery stenosis in mice (CAS)(17). STS treatment (4g/L) decreased IH by about 50% in WT mice (**Figure 1A**), as expressed as the mean intima thickness, the ratio of intima over media thickness (I/M), or the area under the curve of intima thickness calculated over 1 mm. To model the hyperlipidaemic state of patients with PAD, we also performed the CAS on hypercholesterolemic LDLR^-/-^ mice fed for 3 weeks with a cholesterol-rich diet. As expected, the LDLR^-/-^ mice developed more IH than WT mice upon CAS, and STS treatment lowered IH by about 70% (**Figure 1B**). Interestingly, STS did not reduce media thickness in WT mice (p= but significantly reduced media thickness in LDLR^-/-^ mice (p=**Figure 1B**). Of note, the sodium salt H_2_S donor NaHS (0.5g/L) also significantly decreased IH following carotid stenosis in WT mice (**Figure S1 in the online supplementary files)**.

**Figure 1.**
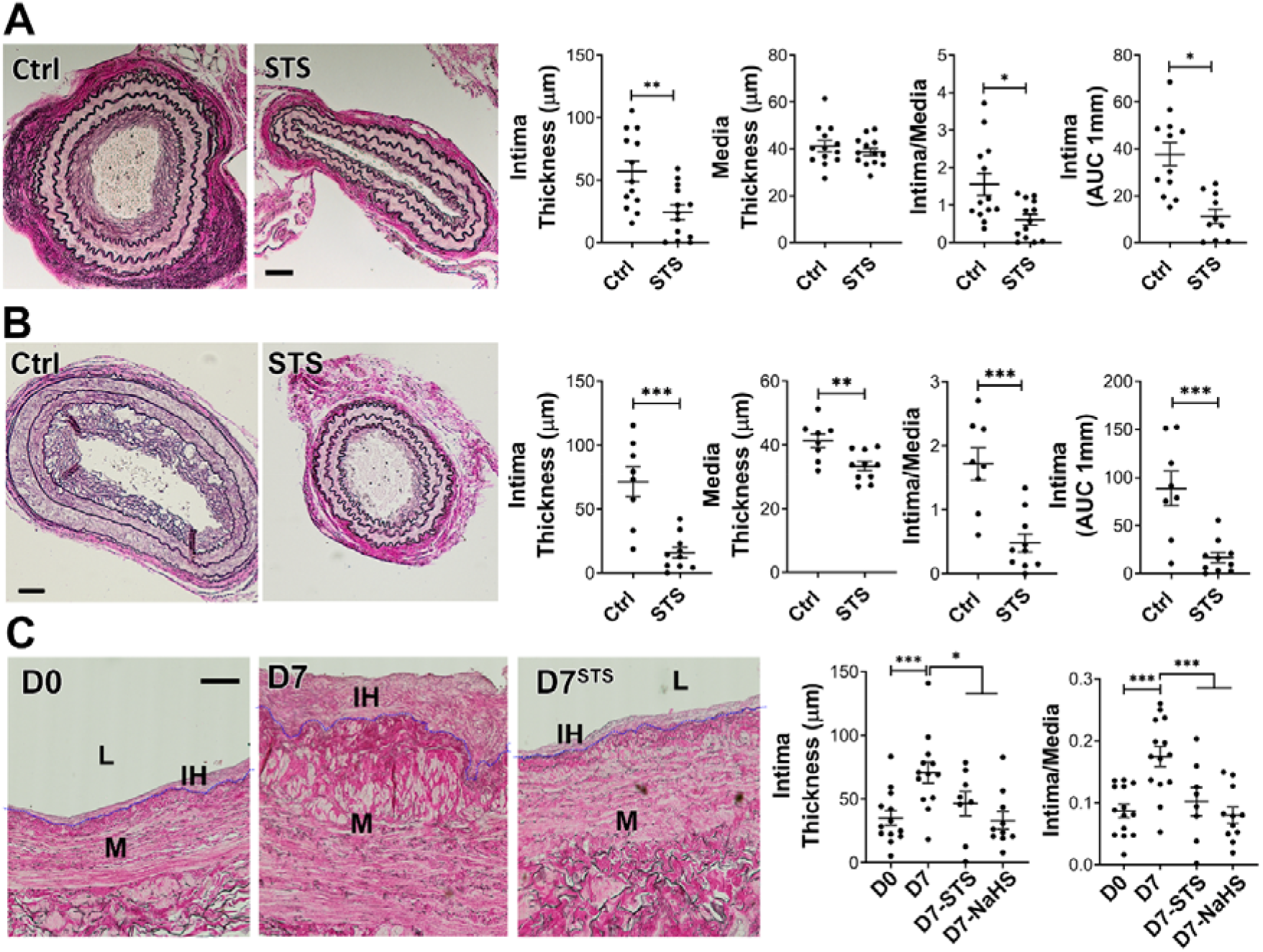
STS and NaHS decrease IH formation after carotid artery stenosis in mice and in cultured human saphenous veins. **A-B**) WT (**A**) or LDLR^-/-^ mice (**B**) treated or not (ctrl) with 4g/L STS were subjected to the carotid artery stenosis surgery. VGEL staining of left carotid cross sections and morphometric measurements of intima thickness, media thickness, intima over media ratio and intima thickness AUC. Scale bar 40µM. Data are mean±SEM of 13 (**A**) and 8 (**B**) animals per group. *p<.05, **p<.01, ***p<.001 as determined by bilateral unpaired t-test. **C**) Intima thickness, media thickness and intima over media ratio of freshly isolated human vein segments (D0) or after 7 days (D7) in static culture with STS (15mM) or NaHS (100µM). Scale bar 60µm. Data are mean±SEM of 12 different veins/patients. *p<0.05, **p<.01, ***p<.001, as determined by repeated measures one-way ANOVA with Dunnett’s multiple comparisons.

### 3.2. STS limits IH development in a model of ex vivo human vein segment culture

Both STS (15mM) and NaHS (100µM) inhibited IH development in our validated model of static *ex vivo* human vein segment culture(12) as measured by intima thickness and I/M ratio (**Figure 1C**). The polysulfide/H_2_S donors diallyl trisulfide (DATS), cysteine-activated donor 5a and GYY4137 also prevented the development of IH in human vein segments (**Figure S2)**.

### 3.3. STS is a biologically active source of sulfur

Overall, STS and “classical” H_2_S donors similarly inhibit IH. To measure whether STS releases detectable amounts of H_2_S or polysulfides, we used the SF_7_-AM and SSP4 probes, respectively. We could not detect any increase in SSP4 or SF_7_-AM fluorescence in presence of STS with or without VSMCs. Na_2_S_3_ was used as a positive control for the SSP4 probe and NaHS as a positive control for the SF_7_-AM probe (**Figure S3)**. The biological activity of H_2_S is mediated by post-translational modification of reactive cysteine residues by persulfidation, which influence protein activity (21, 27). As a proxy for H_2_S release, we assessed global protein persulfidation by DAZ-2-Cy5.5 labelling of persulfide residues in VSMC treated for 4 hours with NaHS or STS. Both STS and NaHS similarly increased persulfidation in VSMC (**Figure 2A**). Using the SSP4 probe, we further observed higher polysulfides levels *in vivo* in the plasma of mice treated for 1 week with 4g/L STS (**Figure 2B**). Similarly, polysulfides levels tended to be higher in the liver of mice treated with STS (p=0.15). As a positive control, mice treated with 0.5g/L NaHS had significantly higher polysulfides levels in the liver (**Figure 2C**). Cse is main enzyme responsible for endogenous H_2_S in the vasculature and Cse^-/-^ mice have been shown to develop more IH (11, 28, 29). Here, we generated a new Cse knock-out mouse line. Consistent with a role as an H_2_S donor, STS fully rescued Cse^-/-^ mice from increased IH in the model of carotid artery stenosis (**Figure 2D**).

**Figure 2.**
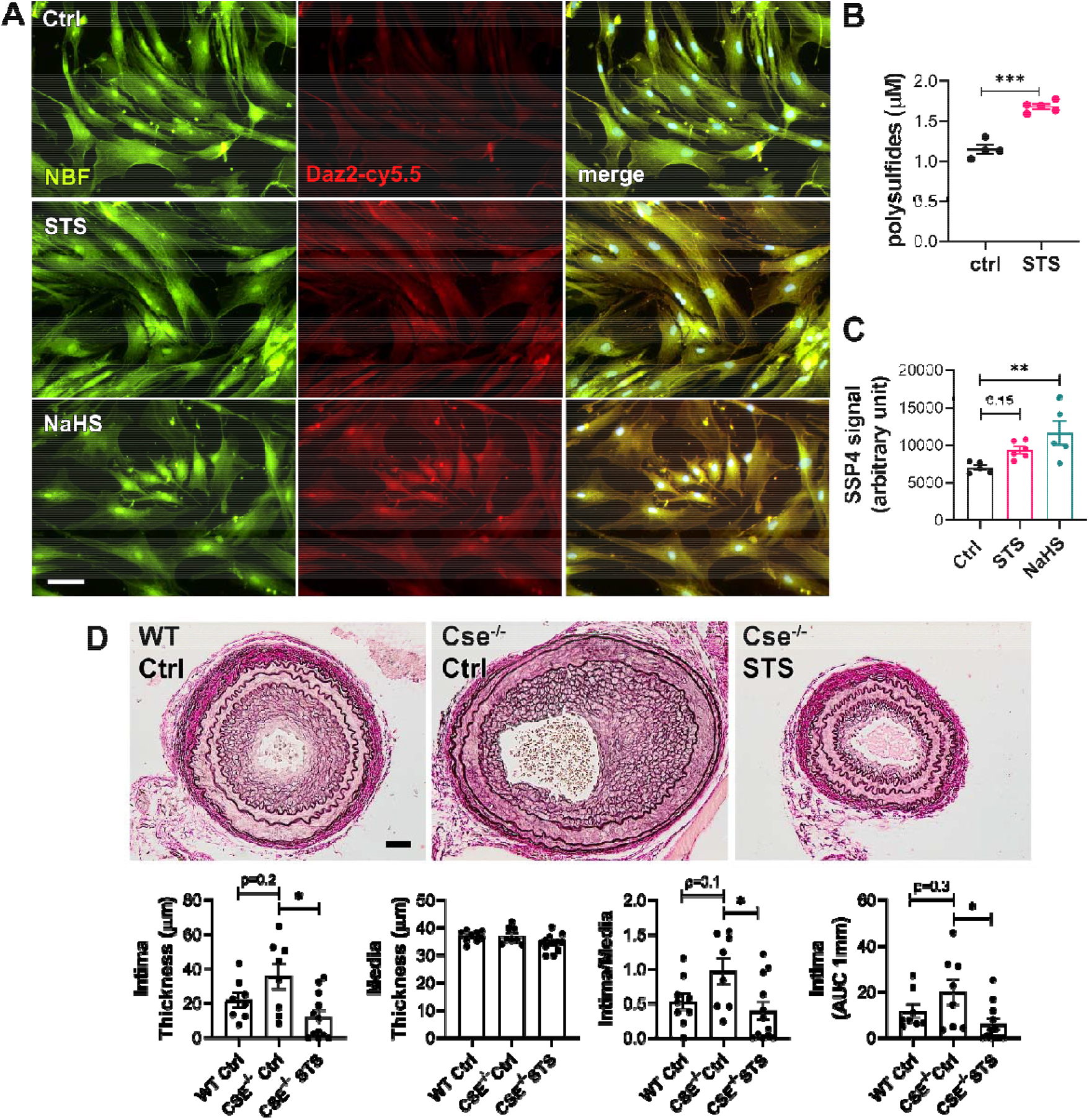
STS increases protein persulfidation. **A**) In situ labeling of intracellular protein persulfidation assessed by DAz-2:Cy5.5 (red), normalized to NBF-adducts fluorescence (green), in VSMC exposed for 4 hours to NaHS (100 µM) or STS (15mM). Representative images of 5 independent experiments. Scale bar 20μm. **B)** Plasma polysulfides levels, as measured by the SSP4 probe, in mice treated 7 days with STS 4g/L. Data are scatter plots with mean ± SEM of 4 animals per group. ***p<.001 as determined by bilateral unpaired t-test. **C)** Polysulfides levels, as measured by the SSP4 probe in liver extracts of mice treated 7 days with STS 4g/L or NaHS 0.5g/L. Data are scatter plots with mean ± SEM of 5 animals per group. **p<.01 as determined by One-way ANOVA with post-hoc t-tests with Tukey’s correction for multiple comparisons. **D)** Intima thickness, media thickness, intima over media ratio and intima thickness AUC of CAS operated mice measured 28 days after surgery in WT mice versus Cse^-/-^ mice treated or not (Cse^-/-^ Ctrl) with 4g/L STS (Cse^-/-^ STS). Scale bar 50µM. Data are scatter plots with mean ± SEM of 8 to 10 animals per group. *p<.05, **p<.01, as determined by one-way ANOVA with post-hoc t-tests with Tukey’s correction for multiple comparisons.

### 3.4. STS limits IH-associated matrix deposition and apoptosis in human vein segments

We further assessed matrix deposition and apoptosis in human vein segments. Concomitant with IH formation, *ex vivo* vein culture (D7) resulted in *de novo* collagen deposition compared to D0, as assessed by polychrome Herovici staining (**Figure 3A)**. Immunoanalysis of immature Collagen III levels confirmed that vein culture (D7) resulted in *de novo* collagen deposition compared to D0, as assessed by Western blot (**Figure 3B)** and immunostaining (**Figure 3C)**. STS and NaHS treatment tended to reduce collagen deposition as assessed by Herovici staining, collagen III immunostaining and Western blotting (**Figure 3A-C)**. TUNEL assay revealed that STS, and to a lesser degree NaHS, decreased the percentage of apoptotic cells observed after 7 days in culture (D7) (**Figure 3D)**. STS also attenuated pro-apoptotic protein Bax overexpression observed after 7 days in culture, while NaHS had a non-significant tendency to decrease Bax level (p=.11; **Figure 3E**). STS also tended to increase the protein level of anti-apoptotic protein Bcl-2 (p=.06), while NaHS significantly increased it (p=.04; **Figure 3E**).

**Figure 3.**
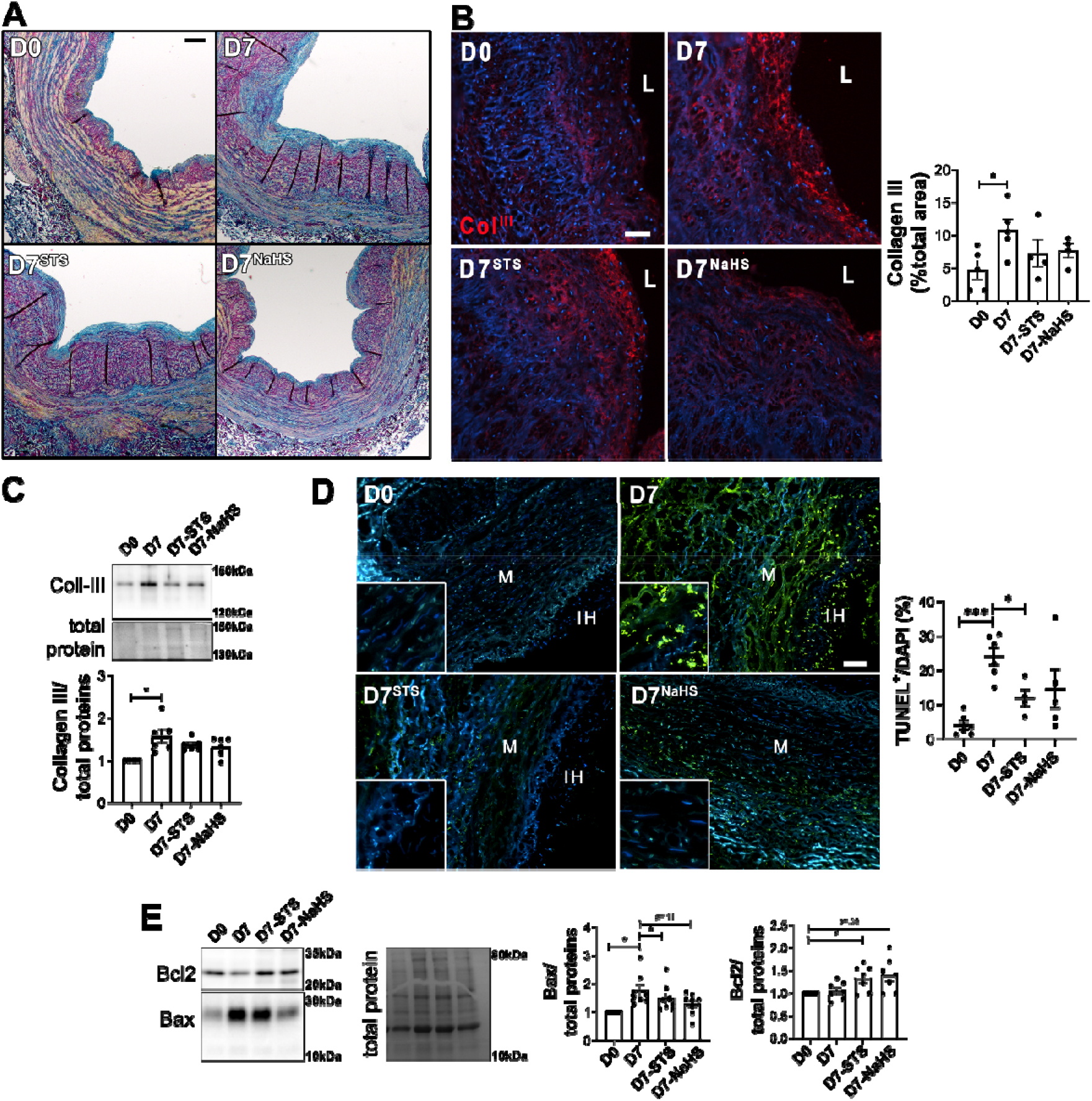
STS decreases apoptosis and matrix deposition in human vein segments. Human vein segment at D0 or after 7 days of static culture with or without (D7) 15mM STS or 100µM NaHS. **A**) Representative Herovici staining of 5 different human vein segments. Mature collagen I is stained pink; new collagen III is stained blue; cytoplasm is counterstained yellow; nuclei are stained blue to black. Scale bar=80µm **B**) *Left panels:* Representative collagen III immunofluorescent staining. Scale bar=50µm. *Right panel* Quantitative assessment of Collagen III immunofluorescent staining. Data are scatter plots of 5 different veins with mean±SEM. *p<0.05 as determined by paired repeated measures one-way ANOVA with Dunnett’s multiple comparisons. **C)** Representative western blot of collagen III over total protein and quantitative assessment of 6 different human vein segments. Data are scatter plots with mean±SEM. *p<0.05 as determined by paired repeated measures one-way ANOVA with Dunnett’s multiple comparisons. **D)** *Left panels:* Representative TUNEL staining in human vein segments. *Right panel:* Apoptosis is expressed as TUNEL positive (green) over DAPI positive nuclei. Scale bar= 50µm. Data are scatter plots of 5 to 6 different veins with mean±SEM. *p<.05, ***p<.001 as determined by mixed model analysis with Dunnett’s multiple comparisons. **E)** Representative western blot of Bax and Bcl2 over total protein and quantitative assessment of 7 different human veins. Data are scatter plots with mean±SEM. *p<0.05 as determined by repeated measures one-way ANOVA with Dunnett’s multiple comparisons.

### 3.5. STS blocks VSMC proliferation and migration

IH is driven by VSMC reprogramming toward a proliferating, migrating and ECM-secreting phenotype (5). Both STS and NaHS significantly reduced the percentage of proliferating cells (defined as PCNA positive nuclei over total nuclei) *in vivo* in CAS-operated carotids in WT mice (**Figure 4A**) and *ex vivo* in human vein segments (**Figure 4B**). *In vitro*, STS dose-dependently decreased primary human VSMC proliferation as assessed by BrdU incorporation assay (**Figure 5A)**. Of Note, NaHS as well as DATS, donor5A and GYY4137, also decreased VSMC proliferation (**Figure S4)**. STS and NaHS also decreased VSMC migration in a wound healing assay (**Figure 5B**). Further evaluation of cell morphology during the wound healing revealed that STS and NaHS-treated cells lost the typical elongated shape of VSMC, as measured through the area, Feret diameter and circularity of the cells (**Figure 5C)**.

**Figure 4.**
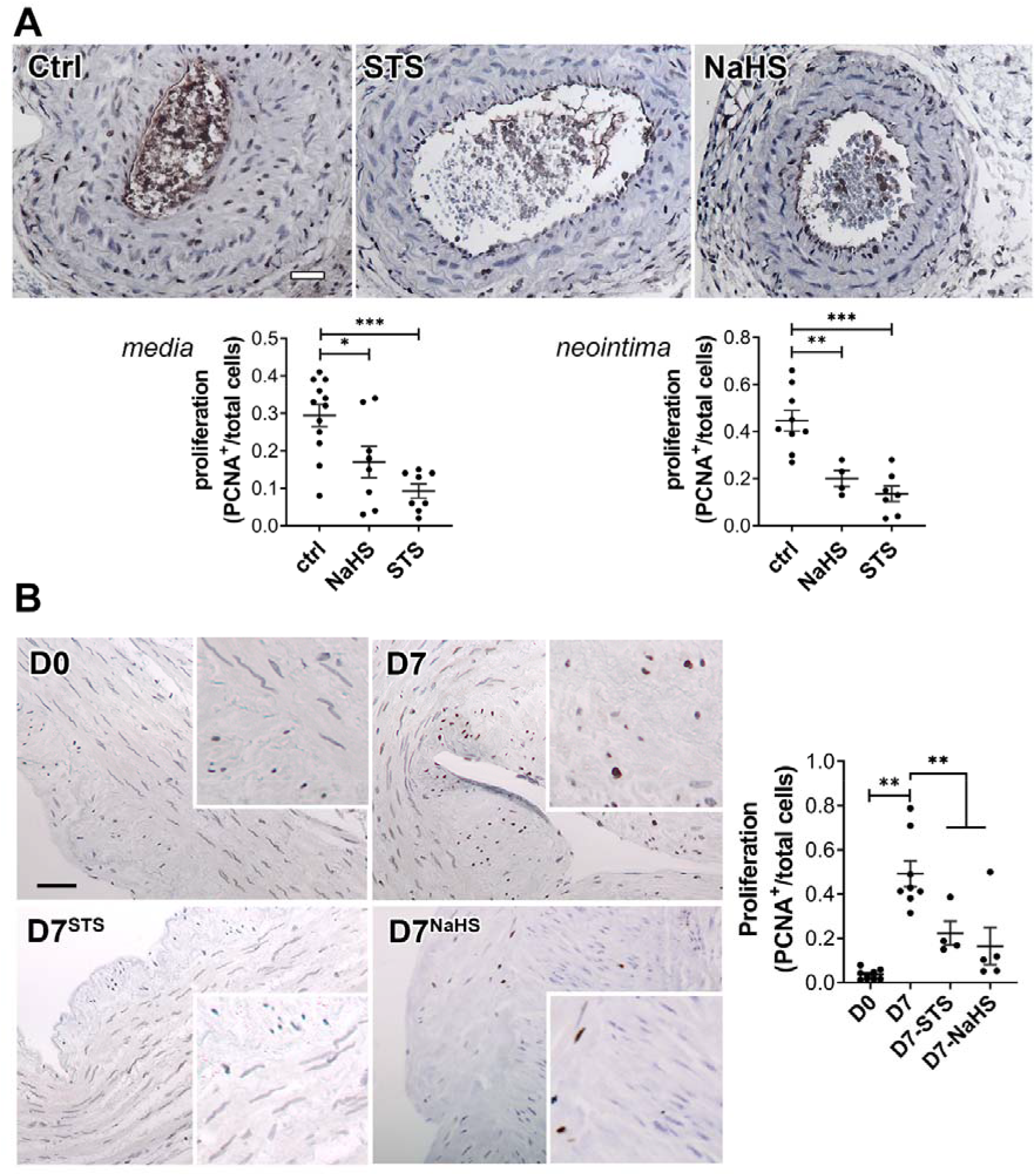
STS inhibits cell proliferation *in vivo* in mouse carotids and *ex-vivo* in human vein segments. PCNA immunostaining on CAS operated carotids in WT mice (**A**) treated or not (Ctrl) with STS 4g/L or NaHS 0.5g/L for 28 days, and human vein segments (**B**) incubated or not (Ctrl) with 15mM STS or 100µM NaHS for 7 days. Proliferation is expressed as the ratio of PCNA positive (brown) nuclei over total number of nuclei. Data are scatter plots with mean±SEM. (**A**) Scale bar 20µm. *p<.05, **p<.01, ***p<.001 as determined from 8 to 12 animals per group by one-way ANOVA with Dunnett’s multiple comparisons. (**B**) Scale bar 50µm **p<.01, as determined from 5 to 7 different veins by mixed model analysis and Dunnett’s multiple comparisons.

**Figure 5.**
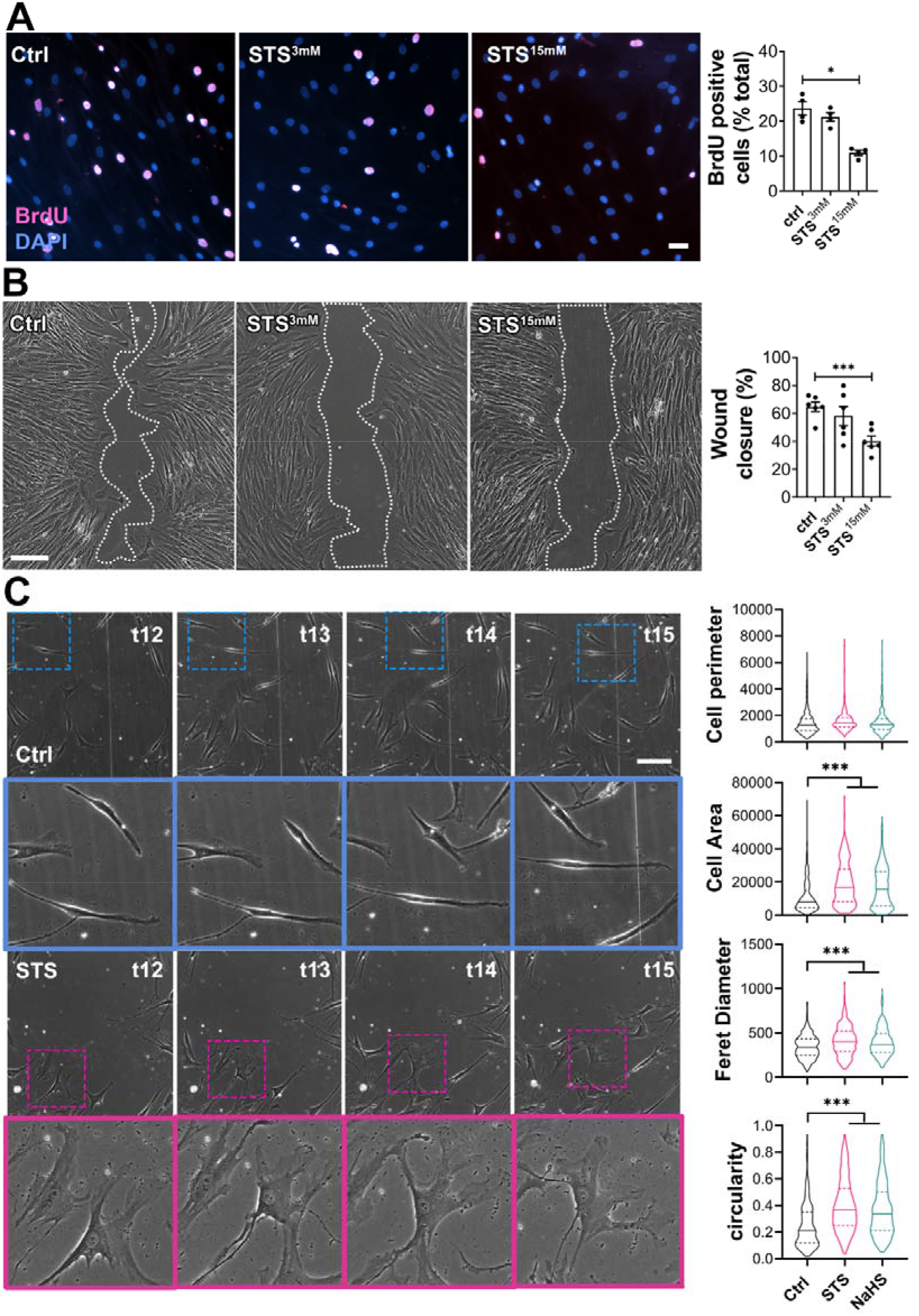
STS inhibits VSMC proliferation and migration *in vitro*. **A**) VSMC proliferation (BrdU incorporation) in cells treated or not for 24 hours with or without (Ctrl) 3 or 15mM STS. Data are % of BrdU positive nuclei (pink) over DAPI positive nuclei. Scale bar: 25µm. Data shown as mean±SEM of 6 different experiments. *p<.05 as determined by repeated measures one-way ANOVA with Dunnett’s multiple comparisons tests. **B)** VSMC migration in cells treated or not for 24 hours with or without (Ctrl) 3 or 15mM STS, as assessed by wound healing assay, expressed as the percentage of wound closure after 10 hours. Scale bar: 100µm. Data are scatter plots with mean±SEM of 5 independent experiments in duplicates. ***p<.001 as determined by repeated measures one-way ANOVA with Dunnett’s multiple comparisons. **C)** Bright field images of VSMC morphology in cells exposed or not (Ctrl) to 15mM STS or 100µM NaHS, as measured as cell perimeter, cell area, Feret diameter and circularity index assessed during wound healing assay. Data are violin plots with median and quartiles (dotted lines) of 5 independent experiments. ***p<.001 as determined by one-way ANOVA with Dunnett’s post-hot test. Scale bar: 80µm. Pink and blue insets are 3-fold magnifications of outlined areas.

### 3.6. STS interferes with microtubules organization

Given the impact of STS on cell morphology, we examined in more details the effect of STS on the cell cytoskeleton. α-tubulin levels were increased in the carotid wall of CAS-operated mice, which were reduced by STS, as demonstrated by immunohistochemistry (**Figure 6A)**. α-tubulin levels were also decreased in the native aorta of mice treated with STS for 7 days (**Figure 6B)**. Similarly, total α-tubulin levels were decreased in *ex vivo* vein segments after 7 days of STS treatment (**Figure 6C-D)**. Looking further at α-tubulin by immunofluorescent staining showed a loss in microtubule in VSMCs treated with STS or NaHS for 8 hours (**Figure 6 E-F)**. To study the effect of H_2_S on microtubule formation, an *in vitro* tubulin polymerization assay was performed in presence of 15mM STS, 100 µM NaHS or 10µM Nocodazole, an inhibitor of microtubule assembly. As expected, Nocodazole slowed down microtubule assembly, as compared to the control. Surprisingly, both NaHS and STS fully blocked microtubule assembly in this assay (**Figure 6G)**. Further studies of the cell cycle in VSMC revealed that 48 hours of treatment with STS or Nocodazole resulted in accumulation of cells in G2/M phase (**Figure 6H)**.

**Figure 6.**
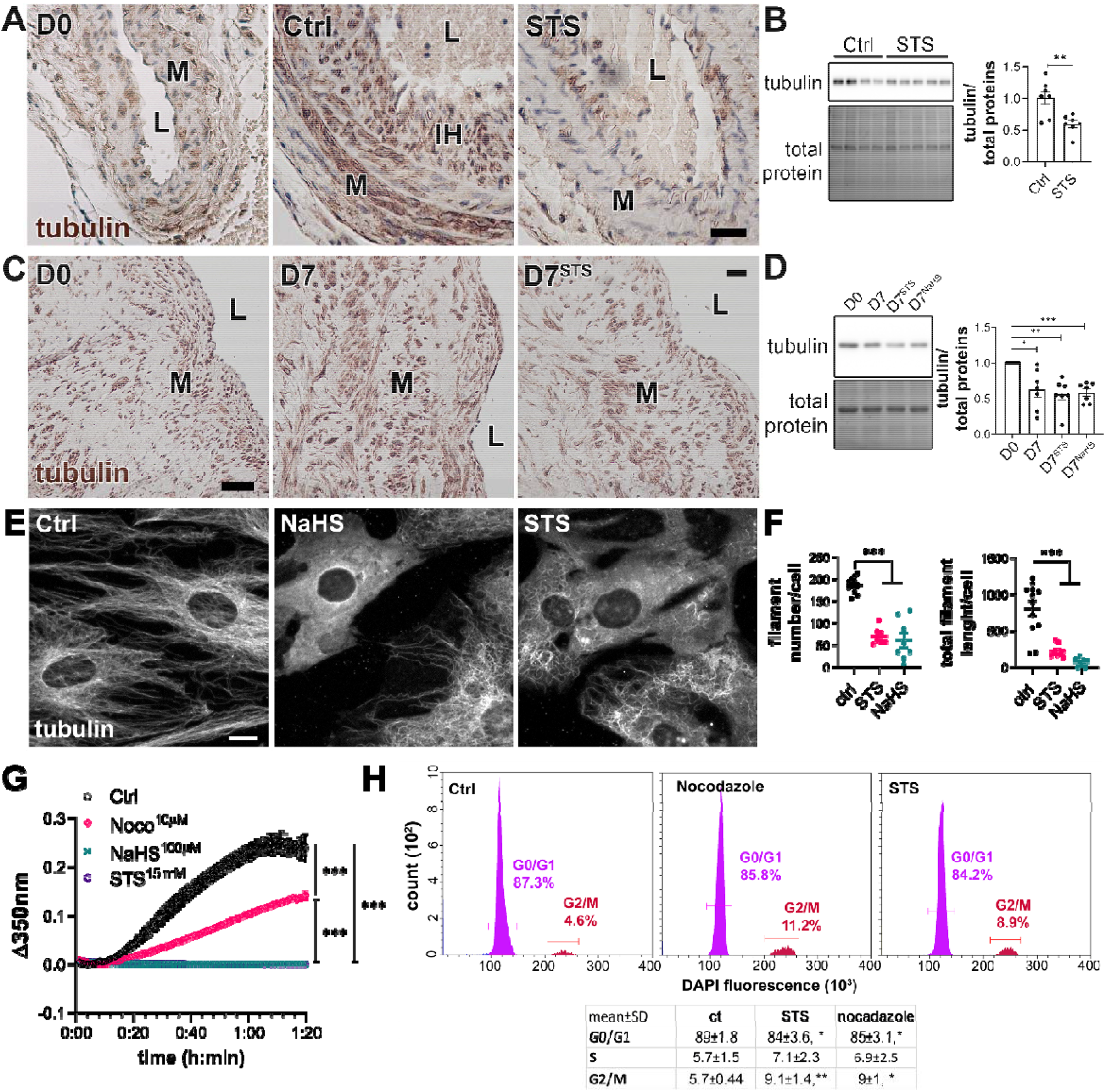
STS inhibits microtubule polymerization in VSMC. **A)** *α*-tubulin immunolabelling in carotids of native (D0) or CAS-operated mice treated or not (Ctrl) with STS 4g/L. L=Lumen; M= Media; IH= Intimal Hyperplasia. Images are representative of 5 to 8 mice per group. **B)** WB analysis of α-tubulin over total protein in aortas of mice treated or not (Ctrl) with STS 4g/L for 7 days. Data are scatter plots of 7 mice per groups with mean±SEM with **p<.01, as determined by one-way ANOVA with Tukey’s multiple comparisons tests. **C)** α-tubulin immunolabelling in human vein segments kept or not (D0) in culture in presence or not (Ctrl) of 15mM STS for 7 days. Scale bar 40µm. L=Lumen; M= Media. Images are representative of 5 different veins. **D)** WB analysis of tubulin over total protein in human vein segments kept or not (D0) in culture in presence or not (Ctrl) of 15mM STS or 100 µM NaHS for 7 days. *p<.05, **p<.01, ***p<.001, as determined by repeated measures one-way ANOVA from 7 different veins with Dunnett’s multiple comparisons tests. **E**) α-tubulin immunofluorescent staining in VSMC exposed or not to 15mM STS or 100 µM NaHS for 8 hours. Images are representative of 5 independent experiments. Bar scale 10 µm. **F)** Quantitative assessment of microtubule filaments immunostaining per cell. Data are representative of 3 independent experiments, 3 to 4 images per experiment per condition. ***p<0.001 as determined by one-way ANOVA with Tukey’s multiple comparisons tests. **G)** *In vitro* tubulin polymerization assay in presence or not (Ctrl) of 15mM STS, 100µM NaHS or 10µM Nocodazole. Data are mean±SEM of 3 independent experiments. **H)** Flow cytometry analysis of cell cycle (DNA content) using DAPI-stained VSMC treated or not (Ctrl) for 48 hours with 15mM STS or 10nM Nocodazole. *Upper panel*: representative histograms; *lower panel*: table with mean±SD of 5 independent experiments. *p<0.05, **p<0.01 as determined by one-way ANOVA with Dunnett’s multiple comparisons tests.

## Discussion

Despite decades of research, intimal hyperplasia remains one of the major limitations in the long-term success of revascularization. In addition, in December 2018, Katsanos and colleagues reported, in a systematic review and meta-analysis, an increased risk of all-cause mortality following application of paclitaxel-coated balloons and stents in the femoropopliteal artery (30). This recent report had tremendous repercussions on the community, urging vascular societies and policy makers to further investigate the use of paclitaxel-coated balloons, stents and other devices for the treatment of peripheral arterial disease. Here, we demonstrate that exogenous sulfur supplementation in the form of STS limits IH development *in vivo* following mouse carotid artery stenosis. Furthermore, STS reduced apoptosis, vessel remodeling and collagen deposition, along with IH development in our *ex vivo* model of IH in human vein segments. We propose that STS limits IH by interfering with the microtubule dynamics, thus VSMC proliferation and migration.

An ever increasing number of studies document the protective effects of H_2_S against cardiovascular diseases (31), including studies showing that H_2_S reduces IH in preclinical models (9-11, 32). The administration of H_2_S in those studies relies on soluble sulfide salts such as NaHS with narrow therapeutic range due to fast and uncontrolled release. Thiosulfate is an intermediate of sulfur metabolism that can lead to the production of H_2_S (15, 33, 34). Importantly, STS is clinically approved and safe in gram quantities in humans. Although STS yields no detectable H_2_S or polysulfide *in vitro*, we observed increased circulating and liver levels of polysulfides in mice, as well as increased protein persulfidation in VSMC. Overall, STS has protective effects against IH similar to the H_2_S salt NaHS and several other “classical” H_2_S donors, but holds much higher translational potential.

Mechanistically, we first observed that STS reduces cell apoptosis and matrix deposition in our *ex vivo* model of human vein segments. This anti-apoptotic effect of STS and NaHS is in line with known anti-apoptotic effects of H_2_S (31). STS also reduces IH *in vivo* following carotid artery stenosis. In this model, fibrosis plays little role in the formation of IH, which relies mostly on VSMC proliferation (5). Therefore, although ECM and especially collagen deposition are major features of IH in human (35, 36), reduced apoptosis and matrix deposition is not sufficient to fully explain the protection afforded by STS in carotids *in vivo*.

STS, similarly to the H_2_S donor NaHS, also inhibits VMSC proliferation and migration in the context of IH *in vivo* in mouse stenotic carotids, as well as *ex vivo* in human vein segments, and *in vitro* in primary human VSMC. These findings are in line with previous studies demonstrating that “classical” H_2_S donors decrease VSMCs proliferation in pre-clinical models (10, 12, 37). In mouse VSMC, exogenous H_2_S has been proposed to promote cell cycle arrest (28), and regulate the MAPK pathway (9) and IGF-1 response (38). Regarding mouse VSMC migration, H_2_S may limit α5β1-integrin and MMP2 expression, preventing migration and ECMs degradation (11, 28).

In this study, we further document that STS and NaHS disrupt the formation of microtubules in human VSMC *in vitro*. Our findings are in line with previous studies showing that Diallyl trisulfide, a polysulfide donor, inhibits microtubule polymerization to block human colon cancer cell proliferation (39). NaHS also depolymerizes microtubules within Aspergillus nidulans biofilms (40). The α/β tubulin dimer has 20 highly conserved cysteine residues, which have been shown to regulate microtubule formation and dynamics (41). In particular, thiol™disulfide exchanges in intrachain disulfide bonds have been proposed to play a key role in microtubule assembly (42). Several high throughput studies of post-translational modification of protein cysteinyl thiols (-SH) to persulfides (-SSH) demonstrated that cysteine residues in α- and β-tubulin are persulfidated in response to H_2_S donors in various cell types (43, 44). Given the prominent role of cytoskeleton dynamics and remodelling during mitosis and cell migration, we propose that STS/H_2_S-driven microtubule depolymerisation, secondary to cysteine persulfidation, contributes to cell cycle arrest and reduces migration in VSMC.

Our findings suggest that STS holds strong translational potential to limit restenosis following vascular surgeries. The dosage of STS used in this study is comparable to previous experimental studies using 0.5 to 2□g/Kg/day (15, 16, 33, 34). In humans, 12.5 and 25□g of STS has been infused without adverse effects (45) and short-term treatment with i.v. STS is safely used in patients for the treatment of calciphylaxis (14). Of note, the pathophysiology of calciphylaxis, also known as calcific uremic arteriopathy (CUA), is caused by oxidative stress and inflammation, which promote endothelial dysfunction, leading to medial remodelling, inflammation, fibrosis and VSMC apoptosis and differentiation into bone forming osteoblast-like cells. Although the main effect of STS on CUA is certainly the formation of highly soluble calcium thiosulfate complexes, our data certainly support the positive effects of STS in the treatment of calciphylaxis. Plus, STS infusions have been shown to increase distal cutaneous blood flow, which could be beneficial in the context of vascular occlusive disease.

Here, we propose that STS treatment results in persulfidation of cysteine residues in the tubulin proteins, which lead to microtubule depolymerization. However, further studies are required to test this hypothesis and demonstrate the STS-induced persulfidation of tubulin cysteine residues. In addition, although we show *in vitro* that H_2_S directly affect microtubule formation in a cell-free environment, other proteins involved in the microtubules dynamics *in vivo* may also be modified by H_2_S, and contribute to the effect of STS on microtubule polymerization and cell proliferation. In this study, STS was administered in the water bottle. However, a validated oral form of the compound has not been developed yet. Case reports and case series suggest that intravenous STS administration is safe, even for relatively long periods of time (46, 47). However, randomized controlled trials testing long-term administration of STS are lacking (47) and the long-term safety and effects of STS administration should be further explored.

In summary, under the conditions of these experiments, STS, a FDA-approved compound, limits IH development *in vivo* in a model of arterial restenosis and *ex vivo* model in human veins. STS most likely acts by increasing H_2_S bioavailability, which inhibits cell apoptosis and matrix deposition, as well as VSMC proliferation and migration via microtubules depolymerization.

## Abbreviations

CAS: carotid artery stenosis
H_2_S: hydrogen sulfide
IH: intimal hyperplasia
NaHS: sodium hydrogen sulfur
PCNA: proliferating cell nuclear antigen
SBP: systolic blood pressure
STS: Sodium Thiosulfate
VSMC: vascular smooth muscle cells
VGEL: Van Gieson elastic lamina

## Contributors

FA, AL and SD designed the study. FA, DM, MMA, ML and SU performed the experiments. FA, DM, MMA, ML and SD analyzed the data. FA, DM, MMA, AL and SD wrote the manuscript. JMC critically revised the manuscript. FA, AL, SD and DM finalized the manuscript.

## Data sharing statement

The data that support the findings of this study are available from the corresponding author, Florent Allagnat, upon request.

## Acknowledgments

We thank the mouse pathology facility for their services in histology (https://www.unil.ch/mpf).

## References

1. Eraso LH, Fukaya E, Mohler ER, 3rd, Xie D, Sha D, Berger JS. Peripheral arterial disease, prevalence and cumulative risk factor profile analysis. European journal of preventive cardiology. 2014;21(6):704–11.

2. Fowkes FG, Rudan D, Rudan I, Aboyans V, Denenberg JO, McDermott MM, et al. Comparison of global estimates of prevalence and risk factors for peripheral artery disease in 2000 and 2010: a systematic review and analysis. Lancet. 2013;382(9901):1329–40.

3. Song P, Rudan D, Zhu Y, Fowkes FJI, Rahimi K, Fowkes FGR, et al. Global, regional, and national prevalence and risk factors for peripheral artery disease in 2015: an updated systematic review and analysis. Lancet Glob Health. 2019;7(8):e1020–e30.

4. Jukema JW, Verschuren JJ, Ahmed TA, Quax PH. Restenosis after PCI. Part 1: pathophysiology and risk factors. Nat Rev Cardiol. 2011;9(1):53–62.

5. Davies MG, Hagen PO. Reprinted article “Pathophysiology of vein graft failure: a review”. Eur J Vasc Endovasc Surg. 2011;42 Suppl 1:S19–29.

6. Zhang L, Wang Y, Li Y, Li L, Xu S, Feng X, et al. Hydrogen Sulfide (H2S)-Releasing Compounds: Therapeutic Potential in Cardiovascular Diseases. Front Pharmacol. 2018;9:1066.

7. Islam KN, Polhemus DJ, Donnarumma E, Brewster LP, Lefer DJ. Hydrogen Sulfide Levels and Nuclear Factor-Erythroid 2-Related Factor 2 (NRF2) Activity Are Attenuated in the Setting of Critical Limb Ischemia (CLI). J Am Heart Assoc. 2015;4(5).

8. Longchamp A, MacArthur MR, Trocha K, Ganahl J, Mann CG, Kip P, et al. Plasma Hydrogen Sulfide Production Capacity is Positively Associated with Post-Operative Survival in Patients Undergoing Surgical Revascularization. 2021:2021.02.16.21251804.

9. Meng QH, Yang G, Yang W, Jiang B, Wu L, Wang R. Protective effect of hydrogen sulfide on balloon injury-induced neointima hyperplasia in rat carotid arteries. Am J Pathol. 2007;170(4):1406–14.

10. Ma B, Liang G, Zhang F, Chen Y, Zhang H. Effect of hydrogen sulfide on restenosis of peripheral arteries after angioplasty. Mol Med Rep. 2012;5(6):1497–502.

11. Yang G, Li H, Tang G, Wu L, Zhao K, Cao Q, et al. Increased neointimal formation in cystathionine gamma-lyase deficient mice: role of hydrogen sulfide in alpha5beta1-integrin and matrix metalloproteinase-2 expression in smooth muscle cells. J Mol Cell Cardiol. 2012;52(3):677–88.

12. Longchamp A, Kaur K, Macabrey D, Dubuis C, Corpataux JM, Deglise S, et al. Hydrogen sulfide-releasing peptide hydrogel limits the development of intimal hyperplasia in human vein segments. Acta Biomater. 2019.

13. Bebarta VS, Brittain M, Chan A, Garrett N, Yoon D, Burney T, et al. Sodium Nitrite and Sodium Thiosulfate Are Effective Against Acute Cyanide Poisoning When Administered by Intramuscular Injection. Ann Emerg Med. 2017;69(6):718–25 e4.

14. Nigwekar SU, Thadhani R, Brandenburg VM. Calciphylaxis. N Engl J Med. 2018;378(18):1704–14.

15. Olson KR, Deleon ER, Gao Y, Hurley K, Sadauskas V, Batz C, et al. Thiosulfate: a readily accessible source of hydrogen sulfide in oxygen sensing. Am J Physiol Regul Integr Comp Physiol. 2013;305(6):R592–603.

16. Snijder PM, Frenay AR, de Boer RA, Pasch A, Hillebrands JL, Leuvenink HG, et al. Exogenous administration of thiosulfate, a donor of hydrogen sulfide, attenuates angiotensin II-induced hypertensive heart disease in rats. Br J Pharmacol. 2015;172(6):1494–504.

17. Tao M, Mauro CR, Yu P, Favreau JT, Nguyen B, Gaudette GR, et al. A simplified murine intimal hyperplasia model founded on a focal carotid stenosis. Am J Pathol. 2013;182(1):277–87.

18. Allagnat F, Dubuis C, Lambelet M, Le Gal L, Alonso F, Corpataux JM, et al. Connexin37 reduces smooth muscle cell proliferation and intimal hyperplasia in a mouse model of carotid artery ligation. Cardiovasc Res. 2017;113(7):805–16.

19. Longchamp A, Alonso F, Dubuis C, Allagnat F, Berard X, Meda P, et al. The use of external mesh reinforcement to reduce intimal hyperplasia and preserve the structure of human saphenous veins. Biomaterials. 2014;35(9):2588–99.

20. Lin VS, Lippert AR, Chang CJ. Cell-trappable fluorescent probes for endogenous hydrogen sulfide signaling and imaging H2O2-dependent H2S production. Proc Natl Acad Sci U S A. 2013;110(18):7131–5.

21. Zivanovic J, Kouroussis E, Kohl JB, Adhikari B, Bursac B, Schott-Roux S, et al. Selective Persulfide Detection Reveals Evolutionarily Conserved Antiaging Effects of S-Sulfhydration. Cell Metab. 2019;30(6):1152–70 e13.

22. Allagnat F, Fukaya M, Nogueira TC, Delaroche D, Welsh N, Marselli L, et al. C/EBP homologous protein contributes to cytokine-induced pro-inflammatory responses and apoptosis in beta-cells. Cell Death Differ. 2012;19(11):1836–46.

23. Teuscher AC, Statzer C, Pantasis S, Bordoli MR, Ewald CY. Assessing Collagen Deposition During Aging in Mammalian Tissue and in Caenorhabditis elegans. Methods Mol Biol. 2019;1944:169–88.

24. Allagnat F, Haefliger JA, Lambelet M, Longchamp A, Berard X, Mazzolai L, et al. Nitric Oxide Deficit Drives Intimal Hyperplasia in Mouse Models of Hypertension. Eur J Vasc Endovasc Surg. 2016;51(5):733–42.

25. Dubuis C, May L, Alonso F, Luca L, Mylonaki I, Meda P, et al. Atorvastatin-loaded hydrogel affects the smooth muscle cells of human veins. The Journal of pharmacology and experimental therapeutics. 2013;347(3):574–81.

26. National Research Council (U.S.). Committee for the Update of the Guide for the Care and Use of Laboratory Animals., Institute for Laboratory Animal Research (U.S.), National Academies Press (U.S.). Guide for the care and use of laboratory animals. 8th ed. Washington, D.C.: National Academies Press; 2011. xxv, 220 p. p.

27. Li Z, Polhemus DJ, Lefer DJ. Evolution of Hydrogen Sulfide Therapeutics to Treat Cardiovascular Disease. Circ Res. 2018;123(5):590–600.

28. Yang G, Wu L, Bryan S, Khaper N, Mani S, Wang R. Cystathionine gamma-lyase deficiency and overproliferation of smooth muscle cells. Cardiovasc Res. 2010;86(3):487–95.

29. Yuan S, Yurdagul A, Jr., Peretik JM, Alfaidi M, Al Yafeai Z, Pardue S, et al. Cystathionine gamma-Lyase Modulates Flow-Dependent Vascular Remodeling. Arterioscler Thromb Vasc Biol. 2018;38(9):2126–36.

30. Katsanos K, Spiliopoulos S, Kitrou P, Krokidis M, Karnabatidis D. Risk of Death Following Application of Paclitaxel-Coated Balloons and Stents in the Femoropopliteal Artery of the Leg: A Systematic Review and Meta-Analysis of Randomized Controlled Trials. J Am Heart Assoc. 2018;7(24):e011245.

31. Szabo C. A timeline of hydrogen sulfide (H2S) research: From environmental toxin to biological mediator. Biochem Pharmacol. 2018;149:5–19.

32. Kip P, Tao M, Trocha KM, MacArthur MR, Peters HAB, Mitchell SJ, et al. Periprocedural Hydrogen Sulfide Therapy Improves Vascular Remodeling and Attenuates Vein Graft Disease. J Am Heart Assoc. 2020;9(22):e016391.

33. Marutani E, Yamada M, Ida T, Tokuda K, Ikeda K, Kai S, et al. Thiosulfate Mediates Cytoprotective Effects of Hydrogen Sulfide Against Neuronal Ischemia. J Am Heart Assoc. 2015;4(11).

34. Lee M, McGeer EG, McGeer PL. Sodium thiosulfate attenuates glial-mediated neuroinflammation in degenerative neurological diseases. J Neuroinflammation. 2016;13:32.

35. Farb A, Kolodgie FD, Hwang JY, Burke AP, Tefera K, Weber DK, et al. Extracellular matrix changes in stented human coronary arteries. Circulation. 2004;110(8):940–7.

36. Suna G, Wojakowski W, Lynch M, Barallobre-Barreiro J, Yin X, Mayr U, et al. Extracellular Matrix Proteomics Reveals Interplay of Aggrecan and Aggrecanases in Vascular Remodeling of Stented Coronary Arteries. Circulation. 2018;137(2):166–83.

37. Yang G, Wu L, Wang R. Pro-apoptotic effect of endogenous H2S on human aorta smooth muscle cells. FASEB J. 2006;20(3):553–5.

38. Shuang T, Fu M, Yang G, Wu L, Wang R. The interaction of IGF-1/IGF-1R and hydrogen sulfide on the proliferation of mouse primary vascular smooth muscle cells. Biochem Pharmacol. 2018;149:143–52.

39. Hosono T, Hosono-Fukao T, Inada K, Tanaka R, Yamada H, Iitsuka Y, et al. Alkenyl group is responsible for the disruption of microtubule network formation in human colon cancer cell line HT-29 cells. Carcinogenesis. 2008;29(7):1400–6.

40. Shukla N, Osmani AH, Osmani SA. Microtubules are reversibly depolymerized in response to changing gaseous microenvironments within Aspergillus nidulans biofilms. Mol Biol Cell. 2017;28(5):634–44.

41. Britto PJ, Knipling L, McPhie P, Wolff J. Thiol-disulphide interchange in tubulin: kinetics and the effect on polymerization. Biochem J. 2005;389(Pt 2):549–58.

42. Chaudhuri AR, Khan IA, Luduena RF. Detection of disulfide bonds in bovine brain tubulin and their role in protein folding and microtubule assembly in vitro: a novel disulfide detection approach. Biochemistry. 2001;40(30):8834–41.

43. Bibli SI, Hu J, Looso M, Weigert A, Ratiu C, Wittig J, et al. Mapping the Endothelial Cell S-Sulfhydrome Highlights the Crucial Role of Integrin Sulfhydration in Vascular Function. Circulation. 2021;143(9):935–48.

44. Fu L, Liu K, He J, Tian C, Yu X, Yang J. Direct Proteomic Mapping of Cysteine Persulfidation. Antioxid Redox Signal. 2020;33(15):1061–76.

45. Farese S, Stauffer E, Kalicki R, Hildebrandt T, Frey BM, Frey FJ, et al. Sodium thiosulfate pharmacokinetics in hemodialysis patients and healthy volunteers. Clin J Am Soc Nephrol. 2011;6(6):1447–55.

46. Brucculeri M, Cheigh J, Bauer G, Serur D. Long-term intravenous sodium thiosulfate in the treatment of a patient with calciphylaxis. Semin Dial. 2005;18(5):431–4.

47. Schlieper G, Brandenburg V, Ketteler M, Floege J. Sodium thiosulfate in the treatment of calcific uremic arteriolopathy. Nat Rev Nephrol. 2009;5(9):539–43.

