## Supplemental data and methods for "Sodium Thiosulfate acts as an H_2_S mimetic to prevent intimal hyperplasia via inhibition of tubulin polymerization"

DATA SUPPLEMENT

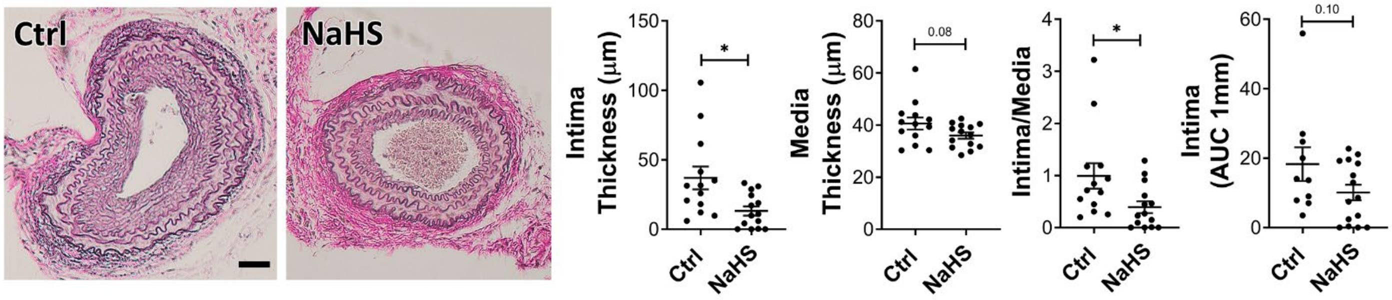


**Figure S1. STS has no effect on systolic blood pressure (SBP) in WT mice.**

WT mice were treated for 28 days post carotid artery surgery with 0.5 gr/L NaHS in the water bottle. Representative VGEL staining of operated left carotid cross sections and morphometric measurements of intima thickness, media thickness, intima over media ratio, and intima thickness AUC over 1mm from the ligation. Data are mean±SEM of 11 to 13 animals per group. *$p<0.05$, as determined by ordinary one-way ANOVA with Dunnett's multiple comparisons.

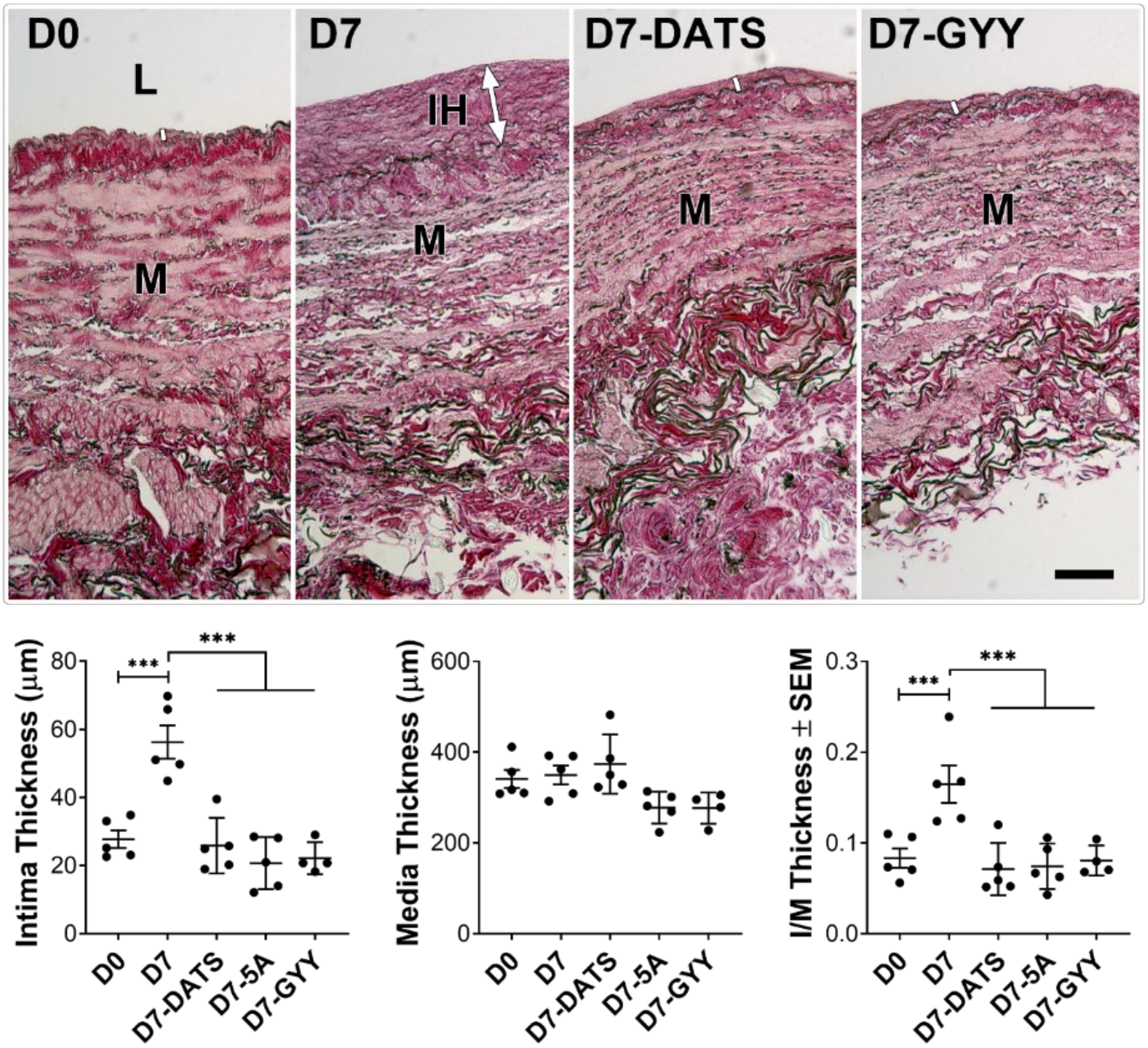


**Figure S2. $H_2S$ donors decrease IH formation in *ex vivo* vein segments.**

VGEL staining of human vein cross section and intima thickness, media thickness and intima over media ratio of freshly collected (D0) human vein segments after 7 days in static culture with or without (D7), diallyl trisulfide (DATS 200 μM), GYY4137 (GYY 200μM) or Donor 5A (30μM). Scale bar: 100 μm. Data are shown as mean±SEM of 5 different veins. *p<.05, **p<.01 as determined by repeated measures one-way ANOVA with Dunnett's multiple comparisons. L=lumen; M= media; IH= intimal hyperplasia

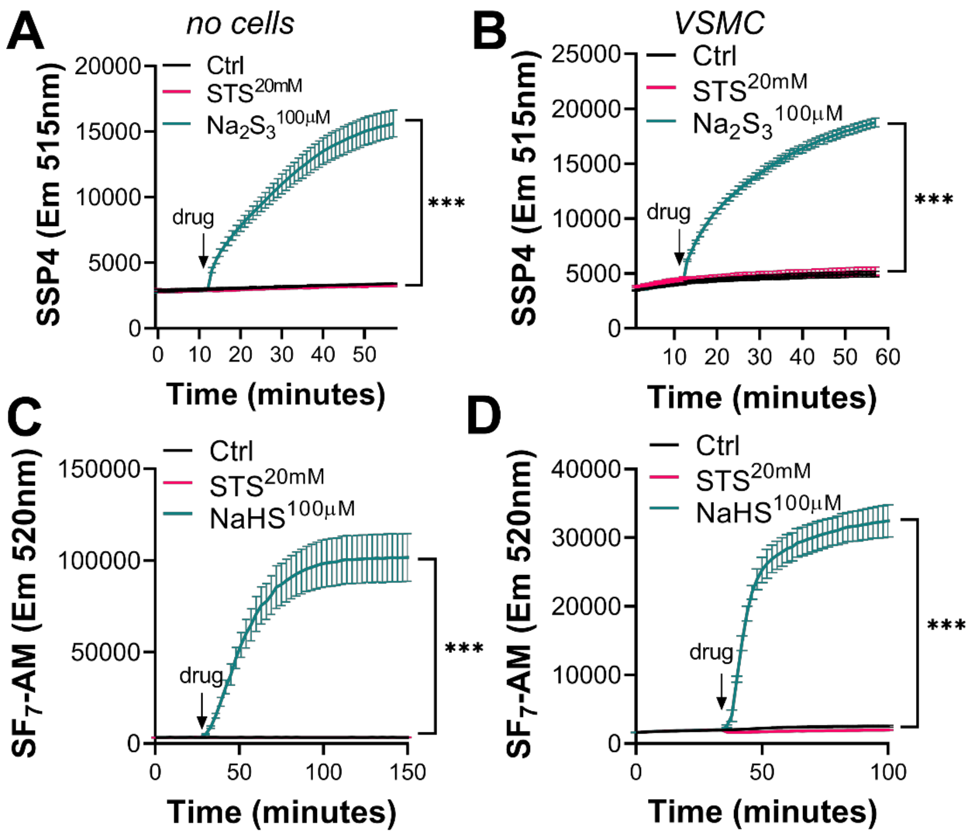


**Figure S3. STS does not release detectable amounts of $H_2S$ or polysulfides *in vitro***

**A-B)** Polysulfides release measured by the SSP4 probe in RPMI media without cells (**A**) or in presence of VSMC (**B**). **C-D)** $H_2S$ release measured by the $SF_7$-AM probe in RPMI media without cells (**C**) or in presence of VSMC **(D)**. Data are mean±SEM of 4 independent experiments. ***$p<.0001$ as determined by repeated measure 2 way ANOVA with Tukey's multiple comparisons.

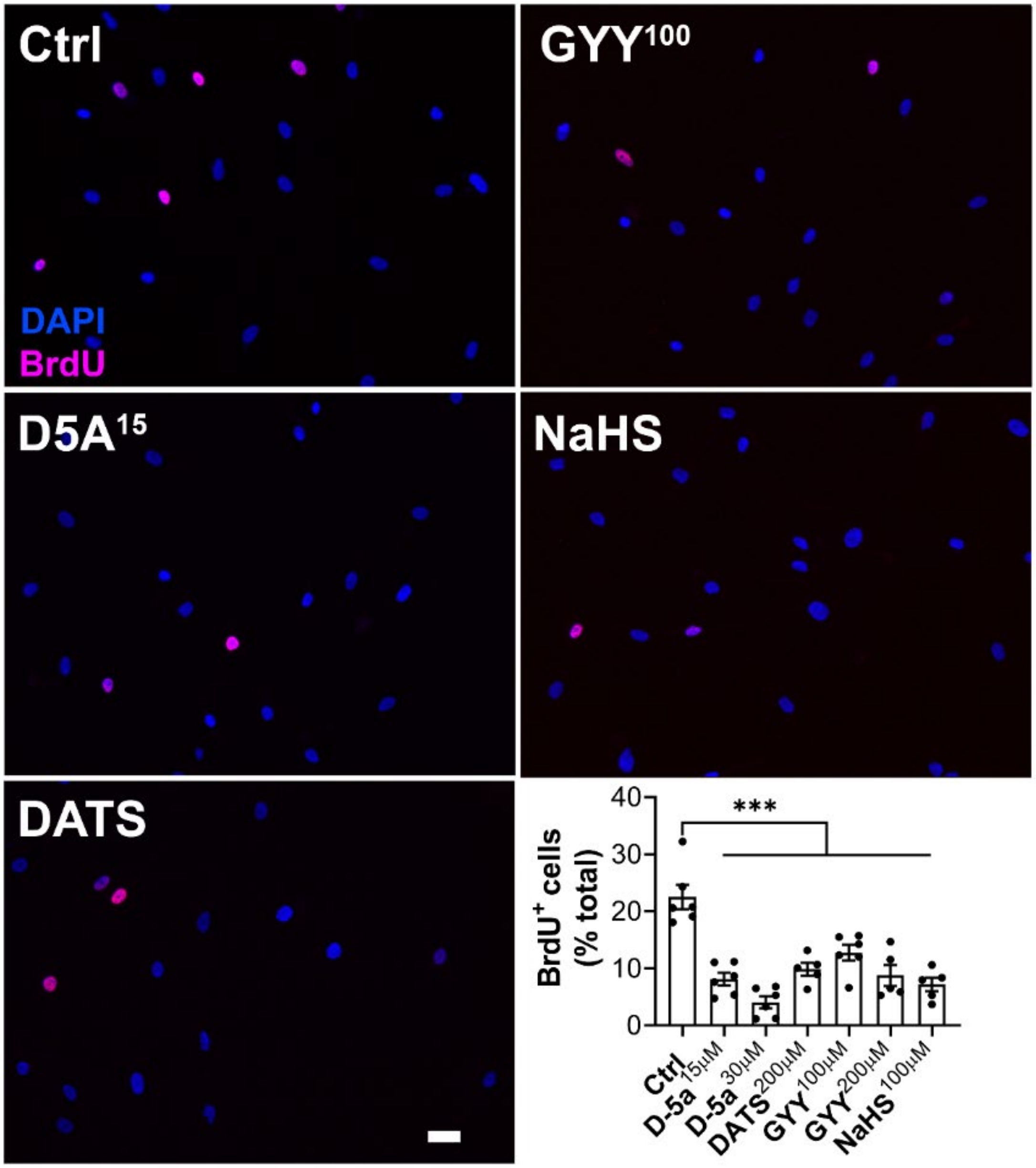


**Figure S4. $H_2S$ donors inhibit VSMC proliferation *in vitro***

VSMC proliferation as assessed by BrdU incorporation (pink) over total nuclei (blue) for 24 hours in presence of absence (Ctrl) of Diallyl trisulfide (DATS 200 µM) or GYY4137 (GYY 100 or 200µM) or Donor 5A (D5A at 15 or 30µM) and NaHS (100µM). Scale bar: 25µm. Data are mean±SEM of 6 independent experiments. ***$p<.001$ as determined by repeated measures ordinary one-way ANOVA with Dunnett's multiple comparisons.

**Supplementary Table S1: Antibodies**

| Target antigen | Vendor | Catalog # | Working concentration | Lot # (preferred but not required) | Persistent ID / URL |
|---|---|---|---|---|---|
| **Collagen III** | Abcam | 7778 | 1/100 | | https://www.abcam.com/collagen-iii-antibody-ab7778.html |
| **Bax** | Santa Cruz | Sc-526 | 1/500 | | https://www.citeab.com/antibodies/807925-sc-526-bax-antibody-p-19 |
| **Bcl2** | Santa Cruz | Sc-492 | 1/500 | | https://www.citeab.com/antibodies/810708-sc-492-bcl-2-antibody-n-19 |
| **Goat anti-Rabbit IgG Secondary Antibody, Alexa Fluor 680** | Thermo Fisher | A21109 | 1/250 | | https://www.thermofisher.com/antibody/product/Goat-anti-Rabbit-IgG-H-L-Highly-Cross-Adsorbed-Secondary-Antibody-Polyclonal/A-21109 |
| **α-Tubulin** | Sigma-Aldrich | T6074 | 1/10000 | | sigma-t6074 |
| **Anti-Rabbit HRPO** | Invitrogen | #31460 | 1/20000 | SJ257972 | Goat-anti-Rabbit-IgG-H-L-Secondary-Antibody-Polyclonal-31460 |
| **Anti-mouse HRPO** | Jackson Immunoresearch | 115-035-146 | 1/15000 | 116724 | Jacksonimmuno-115-035-146 |
| **BrdU** | BD Biosciences | 555627 | 1/200 | 8152642 | bdbiosciences_purified-mouse-anti--brdu_555627 |
| **PCNA** | Dako, Baar, Switzerland | M0879 | 1/100 | 20059892 | proliferating-cell-nuclear-antigen |

**reagents**

| | | |
|---|---|---|
| **Sodium Thiosulfate** | Hänseler AG, Cat:06-6688-01 | |
| **NaHS** | Sigma, Cat:161527 | https://www.sigmaaldrich.com/catalog/product/sigald/161527?lang=fr&region=CH&cm_sp=Insite-_-caSrpResults_srpRecs_srpModel_16721-80-5-_-srpRecs3-1 |
| **DeadEnd Fluorometric TUNEL system** | Promega, G3250 | https://ch.promega.com/products/cell-health-assays/apoptosis-assays/deadend-fluorometric-tunel-system/?catNum=G3250 |
| **Immobilon Western Chemiluminescent HRP Substate** | Millipore, Cat: WBKLS0050 | https://www.merckmillipore.com/CH/de/product/Immobilon-Western-Chemiluminescent-HRP-Substrate,MM_NF-WBKLS0050 |
| **Immobilon-P transfer membrane** | Millipore, Cat: IPVH00010 | https://www.merckmillipore.com/CH/de/product/Immobilon-P-PVDF-Membrane,MM_NF-IPVH00010 |
| **Antifade Mounting Medium with DAPI** | Vectashield, H-1200 | |
| **Pierce reversible protein Stain Kit for PDGF membranes** | Thermo Scientific, Cat: 24585 | https://www.thermofisher.com/order/catalog/product/24585#/24585 |
| **DirectPCR** | Qiagen, 102-T | |
| **Proteinase K** | Qiagen, 1122470 | |
| **RPMI-1640 Glutamax I** | Gibco, Cat: 61870-010 | https://www.thermofisher.com/RPMI |
| **Gelatin type B** | Sigma-Aldrich; Cat: G9391 | G9391 Sigma |

| | | |
|---|---|---|
| **Tripure** | Roche, Cat: 11667157001 | Roche_tripure |
| **DAz-2** | Cayman Chemicals, Cat: 13382 | www.caymanchem.com/product/13382 |
| **Cyanine 5.5 alkyne** | Lumiprobe, Cat: C70B0 | https://www.lumiprobe.com/p/cy55-alkyne |
| **4-Chloro-7-Nitrobenzofurazan** | Sigma Aldrich, Cat: 163260 | https://www.sigmaaldrich.com/catalog/product/aldrich/163260?lang=fr&region=CH |
| **$SF_7$-AM fluorescent probe** | Sigma-Aldrich, Cat: 748110 | Sigmaaldrich_748110 |
| **SSP4 fluorescent probe** | Dojindo Molecular Technologies, SB10 | https://www.dojindo.com/product/sulfobiotics-ssp4-sb10/ |
| **Ketamin** | Ketasol-100, Gräub E.Dr.AG, Bern Switzerland | https://www.graeub.com/fr/products/product/ketasol-100/2054 |
| **Xylasin** | Rompun®, Provet AG, Lyssach, Switzerland | http://www.provet.gr/en/animal-health/products/pharmaceuticals/veterinary-pharmaceuticals/rompun-inj.-sol./7-489 |
| **Buprenorphine** | Temgesic, Reckitt Benckiser AG, Switzerland | Temgesic |
| **Resorcine-Fuchsine Weigert** | Waldeck, Cat: 2E-030 | https://www.reactolab.ch/boutique/chroma/resorcin-fuchsine-weigert-solution-500-ml/ |
| **Hematoxylin crist.** | Merck, Cat: 1.04302.0025 | Hematoxylin-cryst |
| **Acid Fuchsin** | Sigma-Aldrich, Cat: F8129 | https://www.sigmaaldrich.com/catalog/product/sigma/f8129?lang=fr&region=CH |
| **Picric acid** | Sigma-Aldrich, Cat:197378 | https://www.sigmaaldrich.com/catalog/product/aldrich/197378?lang=fr&region=CH |
| **Neo-Clear** | Merck, Cat: 1.09843.5000 | Neo-Clear |
| **Ethanol absolu** | Merck, Cat: 1.00983.2500 | Ethanol |
| **Phosphate Buffered Saline (PBS)** | Bichsel, Cat:100 0 324 | https://www.bichsel.ch/ |
| **SDS** | Promega, Cat: H5114 | https://ch.promega.com/products/biochemicalssodium-dodecyl-sulfate |
| **Tween** | Applichem, Cat: A1389 | https://www.applichem.com/tween |
| **Triton x-100** | Sigma-Aldrich, Cat: T8787 | https://www.sigmaaldrich.com/catalog/product/sigma/t8787?lang=fr&region=CH |
| **BSA** | Applichem, Cat: A1391 | https://www.applichem.comalbumin-fraktion-v |
| **Recombinant Human PDGF-BB** | PeproTech House, Cat: 100-14B | recombinant-human-pdgf-bb |
| ***DC*™ Protein Assay Kit I** | Bio-rad, Cat 5000111 | Bio-rad dc-protein-assay |
